# Non-invasive human skin transcriptome analysis using mRNA in skin surface lipids

**DOI:** 10.1101/2021.04.04.438351

**Authors:** Takayoshi Inoue, Tetsuya Kuwano, Yuya Uehara, Michiko Yano, Naoki Oya, Akira Hachiya, Yoshito Takahashi, Noriyasu Ota, Takatoshi Murase

## Abstract

Non-invasive acquisition of mRNA data from the skin would be extremely useful for understanding skin physiology and diseases. Inspired by the holocrine process, in which the sebaceous glands secrete cell contents into the sebum, we focused on the possible presence of mRNAs in skin surface lipids (SSLs). We found that measurable human mRNAs exist in SSLs, where sebum protects them from degradation by RNases. The AmpliSeq transcriptome analysis was modified to measure SSL-RNAs, and our results revealed that SSL-RNAs predominantly contained mRNAs derived from sebaceous glands, epidermis, and hair follicles. Analysis of SSL-RNAs non-invasively collected from patients with atopic dermatitis revealed significantly increased expression of inflammation-related genes and decreased expression of terminal differentiation-related genes, consistent with the results of previous reports. Further, we found that lipid synthesis-related genes were downregulated in the sebaceous glands of patients with atopic dermatitis. These results indicate that the analysis of SSL-RNAs is promising to understand the pathophysiology of skin diseases.

## Introduction

Intra- and inter-organ communication mediated via various hormones, growth factors, cytokines, metabolites, and miRNAs play important roles in maintaining homeostasis in the human body (1). Several efforts have been made to establish comprehensive analytical methods for these mediators to monitor the physiological conditions of the body and explore predictive biomarkers for various diseases (2–5). Especially, the use of serum, urine, and saliva samples, which can be obtained in a non- or low-invasive manner, has been widely investigated.

The skin is often referred to as “the window to body’s health’’ since the skin phenotypes, such as the cutaneous pathology, appearance, and its secretions reflect not only the skin condition but also the condition inside the body (6). Moreover, the skin forms the body surface and biomolecules can be easily collected from the sweat, hair, and stratum corneum samples, and thus, skin is a useful source of samples to monitor the skin and body conditions. For instance, the cortisol content in the scalp hair correlates with long-term cumulative cortisol exposure (7). The sweat can also be used as an indicator of internal physiological changes (8), and attempts have been made to monitor patients’ conditions, for instance, tracking blood glucose levels by measuring glucose in sweat samples of patients with diabetes (9). Although metabolites, proteins, and DNA are relatively easy to collect from the sweat and hair samples (10, 11), it is difficult to collect measurable mRNAs from skin in a non- or low-invasive manner. So far, tape-stripped stratum corneum has been used to collect mRNAs in a minimally invasive manner (12), however, the mRNA content is very low and highly degraded due to the RNase activity on the skin surface. Therefore, a skin biopsy is practically required to analyze mRNA expression; however, this method is invasive, which limits its application. More recently, a minimally invasive method for mRNA analysis via RNA-seq using AmpliSeq technology with 16–20 consecutive tape strips was reported (13–15). However, tape stripping of the stratum corneum is known to induce skin damage, including disruption of the skin barrier (16), epidermal hyperproliferation, and infiltration of CD3-positive T cells into the dermis (17), indicating that the problems related to the invasiveness of this technique remain to be fully resolved.

The sebaceous glands synthesize and accumulate lipids to produce sebum. The lipids accumulated in the cytosol of sebocytes are secreted into the sebaceous ducts following rupture of the plasma membrane; this mode of secretion is called holocrine secretion (18) and is unique among the exocrine glands such as lipid-secreting sebaceous and meibomian glands. The holocrine secretion of the cell contents led us to the idea that sebum may contain various biomolecules, including mRNAs, which may be useful for analyzing biological information. Therefore, in this study, we first investigated the presence of mRNAs in human sebum and established a non-invasive, comprehensive method of analyzing human mRNAs using skin surface lipids (SSLs) as samples. Further, the applicability of this method in skin characterization was verified in healthy subjects and patients with atopic dermatitis (AD).

## Methods

### Subject recruitment and collection of SSLs

Thirty-two healthy male individuals (mean age: 34.6 years, range: 20–49 years, SD = 9.24) were recruited for the study that was conducted in October 2016. The individuals were evaluated by dermatologists prior to the commencement of the study to confirm no obvious skin disease or condition on their faces. Thirty male patients (mean age: 31.0 years, range: 20–48 years, SD = 8.82) diagnosed with mild or moderate facial AD were recruited during June– October 2017. All subjects were required not to remove facial sebum by washing or using wipes or shaving their face on the test day until the end of the test. Patients with AD were also restricted to use steroidal anti-inflammatory and immunosuppressive drugs on the facial skin 24 h prior to the study. The study was approved by the Human Research Ethics Committee, Kao Corporation (approval numbers: 792-2016082 and T003-170413), Japan Aesthetic Dermatology Symposium (approval number: KU-2017-05-003), and the Shinjukuminamiguchi Dermatologic Clinic (approval number: KU-2016-10-005).

SSLs were collected by wiping the whole face (forehead, cheek, face line, nose, and chin) using an oil blotting film (8.0 cm x 5.0 cm, 3M Japan, Tokyo, Japan) and samples were stored in glass vials at -80 °C until use.

### Skin tissue

Surgically removed adult forehead skin from three Caucasian males, aged 62, 66, and 67 years, and nose skin from one adult (Caucasian female, 66-years-old) were provided by the Colorado Dermatology Institute (Colorado Springs, CO, USA) for laser microdissection and immunostaining, respectively. The procurement of skin tissues was approved by the Institutional Review Board of IntegReview Ltd. (Austin, TX, USA; approval number: T046a-170829) and was conducted according to the Declaration of Helsinki Principles. Informed consents were obtained from the volunteers prior to surgery. After surgery, the skin tissues were stored in William’s E medium (Life Technologies, Carlsbad, CA, USA) at 4 °C until embedding. All skin tissues were embedded using the Tissue-Tek optimal cutting temperature compound (Sakura Finetek, Tokyo, Japan) and kept frozen until sectioning.

### mRNA extraction and qPCR

Human saliva, sweat, urine, and serum samples collected from two donors were purchased from Cosmo Bio (Tokyo, Japan). Total RNA was extracted from 1 mL of each sample using the TRIzol LS reagent (Thermo Fisher Scientific, Waltham, MA, USA). Total RNA was extracted from the stratum corneum of the cheek of two healthy males according to a previous report (12). Collected RNA was dissolved with 10 µL of nuclease-free water. RNAs in SSLs (SSL-RNAs) was extracted using the TRIzol reagent (Thermo Fisher Scientific) as follows: 2.85 mL of TRIzol was added to a finely cut oil blotting film containing sebum samples. Next, the solution was divided equally into two tubes and 260 µL of chloroform was added to each tube and mixed by vortexing. The tubes were centrifuged at 12,000 × *g* for 15 min at 4 °C. The upper layer was transferred to a fresh tube and precipitated with ethanol. The precipitates were washed with 70 % ethanol (v/v) and dissolved in 10 µL of nuclease-free water.

Reverse transcription was performed using the SuperScript IV First-Strand Synthesis System and Oligo-dT primers (Thermo Fisher Scientific). The qPCR was performed using the TaqMan Fast Universal PCR Master Mix (Thermo Fisher Scientific) and TaqMan probes for each gene (Thermo Fisher Scientific).

### Evaluation of mRNA degradation in SSLs

To prepare standard samples with different levels of degraded mRNAs, total RNA (1 µg) extracted from normal human epidermal keratinocytes (NHEK) (Cascade Biologics, Portland, OR, USA) was incubated with 30–1000 ng/mL of recombinant human RNase 7 (Novus Biologicals, Littleton, CO, USA) in 10 mM Tris-HCl buffer (pH 8.0) (Nippon Gene, Tokyo, Japan) for 30 min. QIAzol lysis reagent (Qiagen, Hilden, Germany) and chloroform were added to the samples treated with RNase 7 or SSLs collected from six healthy males as described above. After the solutions were vortexed and centrifuged, RNA in the supernatant was purified using the miRNeasy Mini Kit (Qiagen). The level of RNA degradation in the samples was determined using the High Sensitivity RNA ScreenTape (Agilent Technologies, Palo Alto, CA, USA) on an Agilent 4200 TapeStation system (Agilent Technologies). The level of degradation of human mRNAs in SSLs was estimated using the following calculated DV200 value, which evaluates the percentage of fragments containing > 200 nucleotides. mRNA was reverse-transcribed using the SuperScript IV First-Strand Synthesis System and Oligo-dT primers (Thermo Fisher Scientific) and was amplified in a thermal cycler using the PowerUP SYBR Green Master Mix (Thermo Fisher Scientific) and the following primers:

*ACTB* (forward primer 57 bp): 5’-GCTTTTGGTCTCCCTGGGAG-3’

*ACTB* (forward primer 363 bp): 5’-ACAATGTGGCCGAGGACTTT-3’

*ACTB* (reverse primer): 5’-AGTCAGTGTACAGGTAAGCCC-3’

Abundance of each amplicon was determined from the RNA standard curve. Further, DV200 values of the SSL-RNA samples were calculated from the standard curve of DV200 plotted against abundance ratio of the amplicons (363 bp/57 bp).

### Immunostaining

Skin sections obtained from the nose of a Caucasian female were fixed in acetone at -20 °C for 10 min and treated with 0.1 % Triton X-100/PBS for 5 min. The skin sections were incubated with Protein Block serum-free (Agilent Technologies) for 30 min, then with the primary antibodies against keratin/cytokeratin (mouse mono-clonal (AE-1/AE-3), original solution, Nichirei, Tokyo, Japan) or RNase 7 (rabbit poly-clonal, 1:50, Cloud-Clone Corp., Houston, TX, USA) for 1 h at 20–25 °C, and finally with anti-rabbit IgG (donkey poly-clonal, Alexa Fluor 555, 1:1000, Thermo Fisher Scientific) or anti-mouse IgG (goat poly-clonal, Alexa Fluor 647, 1:1000, Thermo Fisher Scientific) for 30 min. Samples were mounted on a glass slide and imaged using fluorescence microscopy (BZ-X710, Keyence, Osaka, Japan).

### Western blotting

Total protein was extracted from SSLs using 350 μL of RIPA buffer and was purified using the Ready Prep 2-D cleanup kit (Bio-Rad, Hercules, CA, USA); the protein concentration was measured using the BCA protein assay kit (Thermo Fisher Scientific). Afterwards, 70 μL of trichloroacetic acid was added and the samples were incubated on ice for 30 min, followed by centrifugation at 13,000 × *g* for 5 min at 4 °C. Chilled acetone (500 μL) was added to the pellets, the tubes were centrifuged at 13,000 × *g* for 5 min at 4 °C, and the supernatant was removed. The pellets were dissolved in SDS sample buffer (Novagen, Darmstadt, Germany) and boiled at 95 °C for 5 min. Proteins (5 μg) were separated on a 4–15 % polyacrylamide gradient gel (Bio-Rad) and then transferred to a polyvinylidene fluoride (PVDF) membrane (Bio-Rad) soaked in Tris-Glycine Buffer (25 mM Tris, 192 mM glycine; pH 8.2) containing 20 % (v/v) methanol. The PVDF membrane blocked with PVDF Blocking Reagent for Can Get Signal (Toyobo, Tokyo, Japan) was incubated with anti-RNase 7 antibody (rabbit polyclonal, 1:200, Cloud-Clone Corp, Katy, TX, USA) for 60 min followed by incubation with anti-rabbit IgG, horseradish peroxidase-linked secondary antibody (donkey monoclonal, 1:2000, GE Healthcare, Bucks, UK) for 45 min. The bands were visualized using ECL prime western blotting detection reagents (GE Healthcare).

### Effect of sebum lipids on RNase activity

The oil blotting film used to collect SSLs from the face was finely cut and 1 mL distilled water and 4 mL tert-butyl methyl ether were added to the cut film. The solution was transferred to a new glass vial, and centrifuged at 2,050 × *g* at 4 °C for 10 min. The upper layer was transferred to a new glass vial, and the organic solvent was dried by blowing nitrogen over it. The remaining sebum lipids were dissolved in 100 μL dimethyl sulfoxide (DMSO) and used for analysis.

Cholesterol ester (cholesteryl palmitate, Sigma-Aldrich, St. Louis, MO, USA, C6072), wax ester (lauryl palmitoleate, Santa Cruz Biotechnology, Santa Cruz, CA, USA, sc-280908), triacylglycerol (glyceryl trioleate, Sigma-Aldrich, T7140), free fatty acid (palmitoleic acid, Sigma-Aldrich, P9417), squalene (Sigma-Aldrich, S3626), and cholesterol (Sigma-Aldrich, C8667) were used as authentic samples to identify the lipid molecular species. The sebum lipids extracted from the oil blotting films were dissolved in chloroform/methanol (2:1, v/v). The authentic lipids and 5 mg sebum lipid samples were separated on a thin-layer chromatography (TLC) plate using hexane: diethyl ether: acetic acid (70:30:1, v/v/v). After chromatography, the plate was divided using a glass cutter into portions containing the authentic lipids and sebum lipid samples, and the portion containing the authentic lipids was sprayed with a solution containing 10 % (w/v) copper sulfate and 8 % (w/v) phosphoric acid, and then heated at 180 °C for 3 min to visualize the location of each lipid. Based on the mobility of the authentic lipids, the silica corresponding to portions of sebum lipids was scraped off. The silica was sonicated in the chloroform/methanol (2:1, v/v) mixture to extract the lipids and centrifuged at 2,050 × *g* for 5 min. The supernatant was dried and dissolved in 15 μL DMSO; the solution was further diluted 3- and 9-fold using DMSO.

Cholesterol ester (cholesteryl palmitoleate, Olbracht Serdary Research Laboratories, Toronto, Canada, D-161), wax ester (behenyl palmitoleate, Nu Chek Prep, Elysian, MN, USA, WE-1368), triacylglycerol (glyceryl tripalmitoleate, Sigma-Aldrich, T5888; glyceryl trioleate, Sigma-Aldrich, T7140), free fatty acids (myristoleic acid, Sigma-Aldrich, M3525; palmitoleic acid, Sigma-Aldrich, P9417; oleic acid, Sigma-Aldrich, O1008), squalene (Sigma-Aldrich, S3626), and cholesterol (Sigma-Aldrich, C8667) were used to determine the active ingredients in the sebum. NHEK RNA (20 μg/mL final concentration) was mixed with RNase 7 (final concentration: 1 μg/mL) in 10 mM Tris-HCl buffer (pH 8.0) with or without sebum or the lipid reagent (total 50 µL), and sonicated, followed by incubation at 20–25 °C for 30 min. Subsequently, RNA was extracted using TRIzol LS reagent and the quality of the RNA was determined using the High Sensitivity RNA ScreenTape on Agilent 4200 TapeStation System.

### Library preparation for Ion AmpliSeq

Following the addition of 2.85 mL of QIAzol reagent (Qiagen) to a finely cut oil blotting film containing sebum samples, QIAzol solution was divided equally into two tubes. Chloroform (260 µL) was added to each tube and vortexed, and the tubes were centrifuged at 12,000 × *g* for 15 min at 4 °C. The upper layer was transferred to a fresh tube. RNA was purified using the RNeasy mini kit (performing DNase treatment in the purification step) (Qiagen) and eluted from the resin twice using 50 µL of nuclease-free water followed by ethanol precipitation, and then dissolved in 10 µL of nuclease-free water.

To improve the success rate of the library preparation by AmpliSeq protocol, we modified the default protocol of the Ion AmpliSeq Transcriptome Human Gene Expression kit (Thermo Fisher Scientific). Briefly, 1.75 µL RNA solution was mixed with 0.5 µL VILO Reaction Mix and 0.25 µL SuperScript III Enzyme. Reverse transcription was performed at 25 °C for 10 min, 42 °C for 90 min, and finally 85 °C for 5 min. The target DNA amplification was performed by mixing 2.5 µL cDNA solution, 1.5 µL nuclease-free water, 2.0 µL Ion AmpliSeq HiFi Mix, and 4.0 µL Ion AmpliSeq Transcriptome Human Gene Expression Core Panel under the following conditions: 99 °C for 15 sec and 62 °C for 16 min for 20 cycles. The amplified DNA library was purified by mixing 10 µL AMPure XP beads (Beckman Coulter, Miami, FL, USA) according to the manufacturer’s protocol and eluted using 10 µL nuclease-free water. The quality check of the DNA library was conducted using the High Sensitivity D1000 ScreenTape on Agilent 4200 TapeStation. If the DNA library had amplified, a band of approximately 170 bp was observed. After checking the DNA library quality, the reaction solution was prepared by mixing 3.5 µL purified library solution, 2.0 μL Ion AmpliSeq HiFi Mix, 4.0 μL Ion AmpliSeq Transcriptome Human Gene Expression Core Panel, and 0.5 μL VILO Reaction Mix. After adding 1.0 μL FuPa reagent to 10 μL reconstituted reaction solution, the primer sequence was partially digested under the following conditions: 50 °C for 10 min, 55 °C for 10 min, and 60 °C for 20 min. To ligate the adaptor sequence, 2 μL Switch solution, 1 μL Ion Xpress Barcode adapters, and 1 μL DNA ligase were added to 11 μL reaction solution, followed by incubation at 22 °C for 60 min and 72 °C for 5 min. The library (15 μL) ligated with the adaptor sequence was purified via mixing with 18 μL AMPure XP beads according to the manufacturer’s protocol. Libraries were eluted using 50 µL Library Amp Mix (Thermo Fisher Scientific) to which, 2 µL Library Amp Primers were added. The library amplification was conducted using following steps: 98 °C for 15 seconds and 64 °C for 1 min for five cycles. Next, 50 µL PCR product was mixed with 25 µL AMPure XP beads and the supernatant was transferred to fresh PCR tubes. The supernatants were mixed with 60 µL AMPure XP beads and purified; target fragments were eluted from the beads using 10 µL TE buffer. The quality check of the library was performed using the High Sensitivity D1000 ScreenTape on Agilent 4200 TapeStation.

### Sequencing

The library was quantified using the Ion Library TaqManTM Quantitation Kit (Thermo Fisher Scientific). After an input of 50 pM DNA library in the Ion Chef System (Thermo Fisher Scientific), template preparation and chip loading were performed, and RNA-seq was conducted on the Ion S5 XL System (Thermo Fisher Scientific).

### Verification of the correlation coefficient between AmpliSeq and qPCR results

For identifying *RPLP0*, *CDSN*, and *CCL17* expression in SSL-RNAs, cDNA was pre-amplified in 14 cycles using the TaqMan PreAmp Master Mix (Thermo Fisher Scientific) and pooled TaqMan probe (*RPLP0, CDSN*, and *CCL17*) (Thermo Fisher Scientific) and then diluted 5-fold with nuclease-free water. The qPCR was performed according to the aforementioned “mRNA extraction and qPCR” method. Expression value of *RPLP0* was used as an internal control. Correlation between the value of reads per million mapped reads (RPM) of AmpliSeq and the relative expression value of qPCR in healthy subjects and patients with AD was analyzed.

### Laser microdissection (LMD) and AmpliSeq transcriptome analysis

Frozen skin sections (thickness: 10 µm) from three Caucasian males were mounted on membrane slides (PEN-Membrane 2.0 µm, Leica Microsystems, Wetzlar, Germany) treated with 0.1 % (w/v) poly-L-lysine (Fujifilm Wako Pure Chemical, Osaka, Japan). In addition, two frozen sections were directly collected into 750 µL RLT buffer (Qiagen) containing 40 mM dithiothreitol (Sigma-Aldrich) to analyze the transcriptome of the whole tissue. The LMD sections were fixed with acetone at -20 °C for 10 min, stained with 0.05 % (v/v) toluidine blue, and finally dried. Epidermis, sebaceous glands, sweat glands, hair follicles, and dermis were carefully microdissected from the skin sections using LMD7000 (Leica Microsystems). Fifteen target regions from three skin tissues were dissected and dissolved in 50 µL RLT buffer containing 40 mM dithiothreitol. Total RNA was extracted and purified using the RNeasy mini kit (performing DNase treatment in the purification step) (Qiagen). The RNA was concentrated via ethanol precipitation and dissolved in 10 µL nuclease-free water. The RNA quality was checked on the 4200 TapeStation System and cDNA library was prepared for AmpliSeq transcriptome sequencing using 75 pg total RNA according to the method described in “Library preparation for Ion AmpliSeq”. The number of target amplification was changed from 20 to 18 cycles except for dermis samples.

### Statistics, normalization, and differential expression analysis of the AmpliSeq whole transcriptome

All statistical analyses and normalization of the RNA-seq transcriptome data were performed using the R statistical language. Read counts were generated using the AmpliSeq RNA plugin in Ion Torrent Suite Software (Thermo Fisher Scientific) and normalized with the DESeq2 R package (Bioconductor). For differential expression analysis between healthy subjects and patients with AD, a likelihood ratio test was performed with DESeq2 using normalized counts. Heat maps were generated using the heatmap3 package. Dimensionality reduction using t-distributed stochastic neighbor embedding (t-SNE) was performed using the Rtsne function of the Rtsne package. All plots were generated using the tidyverse package in combination with the reshape2, gplots, ggplot2, grid, and cowplot packages.

## Results

### Measurable human mRNA is present in SSLs

qPCR was conducted to evaluate the mRNA abundance in SSL samples, stratum corneum, urine, serum, saliva, and sweat samples. The expression of *ACTB* and *GAPDH* mRNA in SSLs was comparable to their expression in 100 ng–500 pg and 1 ng–100 ng total RNA in NHEK, respectively, whereas these transcripts were largely undetectable in the other body fluid samples (Fig. 1a). The level of human mRNA degradation in SSLs could not be measured directly due to the presence of bacterial mRNA. Therefore, we established an assay to verify the quality of human mRNA in SSLs. The reverse primer was designed near the 3’ end and was used with the forward primers to amplify 363 bp and 57 bp long human *ACTB* mRNA. After reverse transcription using oligo-dT primers, *ACTB* levels were quantified by performing qPCR using primers generating 363 bp and 57 bp human *ACTB* mRNA. In the case of *ACTB* mRNA of longer than 363 bp, both 363 and 57 bp fragments were amplified, whereas in case of *ACTB* mRNA of 57–363 bp, only the 57 bp fragment was amplified (Fig. 1b). This assay was validated using RNA subjected to artificial and gradual degradation using RNase. The gradually degraded RNA was analyzed to calculate the percentage of fragments containing > 200 nucleotides (DV200 value) (Fig. 1c, d). The DV200 values of gradually degraded RNAs negatively correlated with the abundance ratio of the 363 bp and 57 bp amplicons (Fig. 1e). The level of human mRNA degradation in SSLs collected from six males was calculated using a standard curve (Fig. 1e) and showed a mean DV200 value of 56.5 % (Fig. 1f).

**Figure 1.**
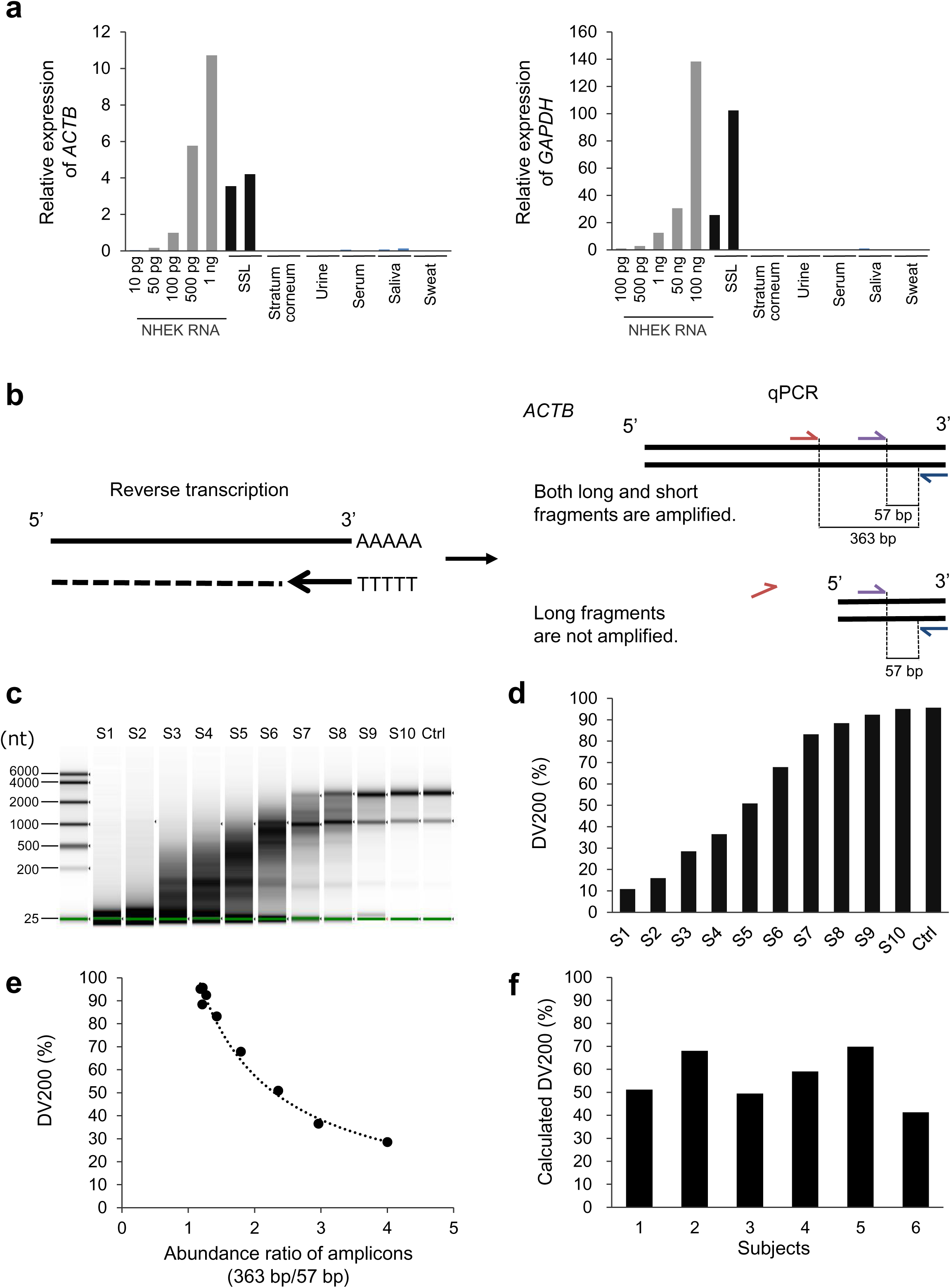
Evaluation of mRNA expression and RNA degradation in skin surface lipids (SSLs) **(a)** Expression of *ACTB* and *GAPDH* mRNA in SSLs, stratum corneum, urine, serum, saliva, and sweat samples analyzed by qPCR. Gene expression is shown as the relative expression for 100 pg of RNA derived from normal human epidermal keratinocytes (NHEK). NHEK total RNA (10 pg to 100 ng) is the standard used. **(b)** Outline for assaying the degradation of human mRNA in SSLs. The extent of mRNA degradation was determined using the long (363 bp)/short (57 bp) amplicon ratio calculated from qPCR results. **(c)** Preparation of RNA samples with different levels of degradation. A series of standards (S1 to S10) are presented with known level of RNA degradation. **(d)** DV200 value indicating the percentage of fragments containing > 200 nucleotides, of 1**c**. **(e)** The relationship between the DV200 of 1**d** and the value of 363 bp/57 bp (abundance of 363 bp amplicon/abundance of 57 bp amplicon measured by qPCR). **(f)** The indirect assessment of the human mRNA degradation in SSLs, n = 6. DV200 of each SSL-RNA sample was calculated using the standard curve of 1**e**.

### Sebum lipids inhibit RNase activity

Consistent with the results of a study reporting RNase 7 expression in human skin (19), we confirmed that RNase 7 was expressed in the sebaceous glands and epidermis and also detected in SSLs (Fig. 2a, b). Because there is high abundance of RNase on the surface of human skin, it was surprising to detect human mRNA in SSLs. This finding led us to hypothesize that sebum lipids inhibit RNase activity. Based on these results, we evaluated the influence of sebum lipids on recombinant RNase 7 activity. Intact cellular RNA and RNase 7 were incubated with or without sebum lipids from four subjects at 37 °C for 30 min. Interestingly, the 28S and 18S ribosomal RNAs were completely degraded after incubation with RNase 7 in the absence of sebum lipids; on the other hand, ribosomal RNAs were stable in the presence of lipids (Fig. 2c). Next, we aimed to identify the key lipids that inhibit RNase 7 activity. The sebum lipids were separated into fractions A–D on a TLC plate (Fig. 2d), and the lipids were recovered from each fraction and subjected to the RNase 7 inhibition assay. We observed that while fractions A, B, and C decreased RNase 7 activity, the inhibitory effect of the fraction D was weak (Fig. 2e). Triglycerides and esters in the sebum are hydrolyzed by the skin microbiome to generate free fatty acids (FFAs). The FFAs in human sebum are predominantly composed of 16 carbon atoms (palmitic acid, 16:0; sapienic acid, 16:1Δ6; and palmitoleic acid, C16:1Δ9) (14). Therefore, we evaluated the inhibitory effects of free palmitoleic acids and various lipids with palmitoleic acid as their main fatty acid, on RNase activity. The free palmitoleic acids and other lipids was dissolved in the reaction buffer at 1 mg/mL or 100 mg/mL concentration, except cholesterol and wax esters, and cholesterol, which could not be dissolved at 100 mg/mL. Our results showed that FFAs strongly suppressed RNase activity at 1 mg/mL compared to other lipids (Fig. 2f). Moreover, FFAs of different chain lengths (myristoleic acid (C14:1) and oleic acid (C18:1)) also suppressed RNase 7 activity (Fig. 2g).

**Figure 2.**
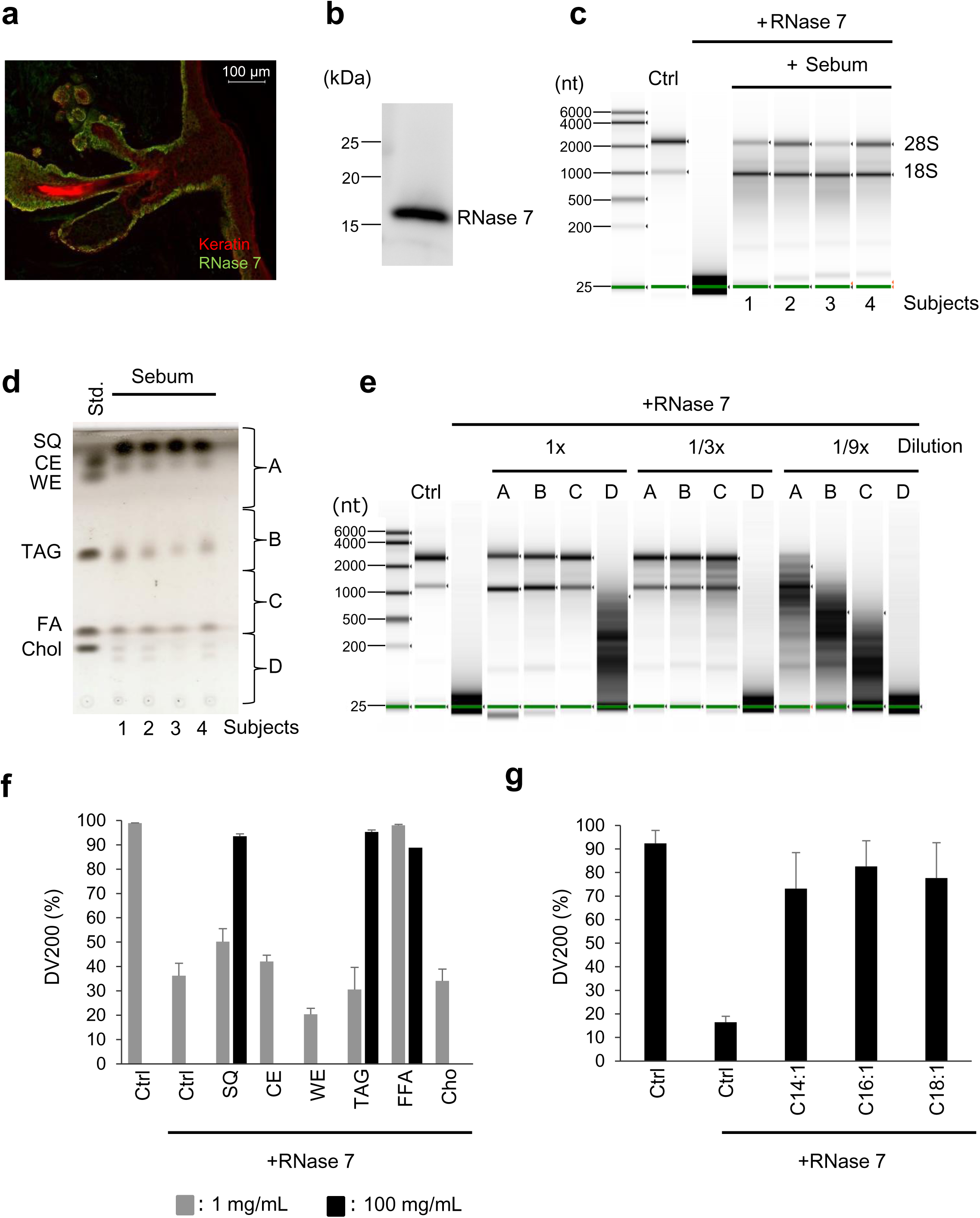
Effect of sebum lipids on RNase activity **(a)** The localization of RNase 7 in the human skin. Green, RNase 7; red, keratin/cytokeratin. Bar: 100 μm. **(b)** The detection of RNase 7 in SSLs collected from healthy males by western blotting. **(c)** Determination of the effect of sebum lipids collected from four healthy males on RNase activity using NHEK total RNA. **(d)** Fractionation of sebum lipids collected from four healthy males by performing thin-layer chromatography (TLC). The standard lane (Std) includes authentic samples: SQ, squalene; CE, cholesterol ester (cholesteryl palmitate); WE, wax ester (lauryl palmitoleate); TAG, triacylglycerol (glyceryl trioleate); FA, free fatty acid (palmitoleic acid); and Chol, cholesterol. A to D sebum samples were collected, and lipids were extracted from the silica gel for the subsequent assays. **(e)** Effect of pooled sebum lipids collected from A to D on RNase activity using NHEK RNA. **(f)** Effect of each lipid on RNase activity using NHEK RNA. The DV200 values are shown as mean ± SE, n = 3. WE, wax ester (behenyl palmitoleate); TAG, triacylglycerol (glyceryl tripalmitoleate); Ctrl; EtOH. **(g)** Relationship between RNase activity and chain length of fatty acids. The DV200 values are shown as the mean ± SE, n = 6. C14:1, myristoleic acid; C16:1, palmitoleic acid; C18:1, oleic acid; Ctrl; EtOH.

### Global expression analysis of SSL-RNAs

For specific and comprehensive quantification of human mRNAs in SSLs, we performed the AmpliSeq transcriptome analysis that can perform multiplexed amplification of cDNA amplicons for more than 20,000 genes. Moreover, this method can analyze even small amounts of RNAs as well as degraded RNAs (20, 21). Although we attempted to prepare sequence libraries from SSL-RNAs based on the default protocol, our success rate was low. Since the data quality can be improved via optimization of the AmpliSeq protocol (22), we modified the experimental conditions for preparing the sequence library. In the modified protocol, we altered the volume of reagents, standardized the conditions for reverse transcription and target amplification, and added a purification step after the target amplification to remove the primer dimers (Supplementary Method 1). With the new protocol, the success rate of the library preparation improved significantly and the AmpliSeq library was prepared with samples obtained from healthy subjects (91 %, 29/32) and patients with AD (100 %, 30/30). To analyze the experimental bias of our protocol, we verified the correlation between the expression results of AmpliSeq and qPCR of thymus and activation-regulated chemokine (*TARC*/*CCL17*) and corneodesmosin (*CDSN*), and observed a high correlation (*CCL17*, R = 0.81, *CDSN*, R = 0.87) (Fig. 3a). Furthermore, the correlation coefficients for the technical replicate of the reverse transcription (0.94 and 0.90) confirmed that our protocol had low experimental bias (Fig. 3b).

**Figure 3.**
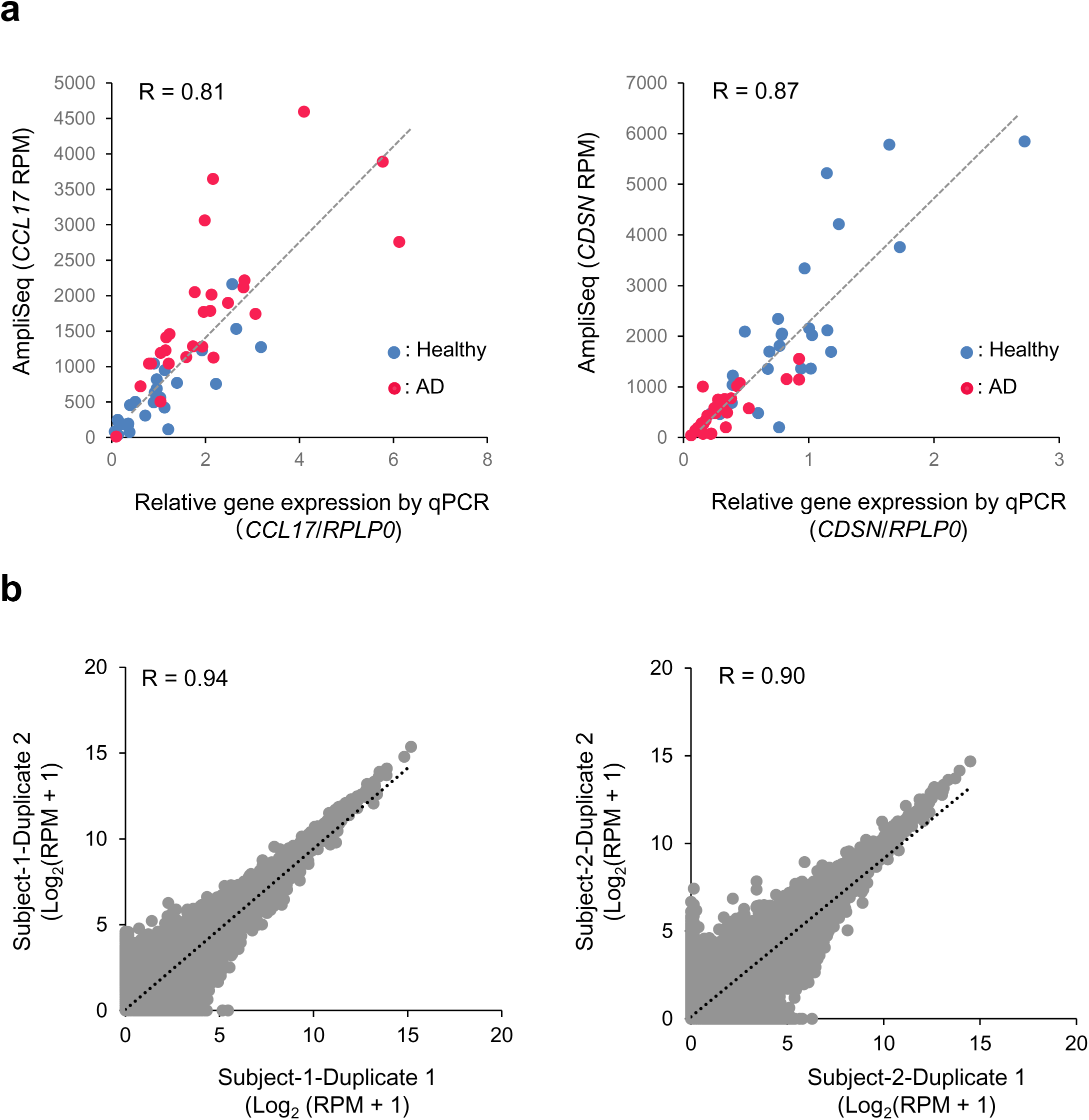
Accuracy of the AmpliSeq data output using the modified protocol **(a)** The correlation between AmpliSeq and qPCR results for *CCL17* and *CDSN* expression in healthy subjects (n = 29, blue) and patients with AD (n = 30, red). **(b)** The correlation of the transcriptome profile when preparing libraries in duplicate from each SSL-RNA obtained from two healthy male subjects.

### The SSL-RNA expression profile predominantly reflects mRNA expression in sebaceous glands, epidermis, and hair follicles

The regions of the sebaceous glands, epidermis, sweat glands, hair follicles, and dermis were isolated from the human skin sections using LMD followed by AmpliSeq transcriptome analysis (Supplementary Fig. 1a). Each region generated distinct clusters when multidimensional scaling (MDS) was performed to analyze the similarity of the transcriptome profile in different regions (Supplementary Fig. 1b). Next, we focused on the genes highly expressed in each region (Supplementary Fig. 1c) (23–38). Genes encoding ELOVL fatty acid elongase 3 and 5 (*ELOVL3* and *ELOVL5*), perilipin 2 and 5 (*PLIN2* and *PLIN5*), and microsomal glutathione S-transferase 1 (*MGST1*) are highly expressed in sebaceous glands (23–26). In our study, these genes were highly expressed in sebaceous glands isolated via LMD than in other regions (Supplementary Fig. 1c). Other regions isolated by LMD also expressed region- characteristic genes, indicating that the LMD was performed successfully.

To gain further insight into the characteristics and origin of SSL-RNAs, we explored genes highly expressed in each region isolated by LMD. In each region, genes with a mean of log_2_ (normalized counts + 1) > 10, and with more than 1.5-fold differential expression compared to other region(s), were selected (Fig. 4). However, using these criteria, we did not find any gene selectively expressed in epidermis, due to its similarity with the hair follicles. Therefore, we listed epidermal genes with more than 1.5-fold differential expression than in sebaceous glands, sweat glands, and dermis. To obtain information regarding the origin of SSL-RNAs, the SSL-RNA profile of 29 healthy subjects was analyzed using the genes expressed characteristically at each region. The results presented in the heat map indicate that SSL-RNAs predominantly comprise mRNAs characteristic for sebaceous glands, epidermis, and hair follicles (Fig. 4). The filaggrin (*FLG*), filaggrin 2 (*FLG2*) and aspartic peptidase retroviral like 1 (*ASPRV1*) that were expressed in the granular layer of epidermis (27–29), were abundantly expressed in SSL-RNAs (Fig. 4). The genes encoding keratin 25, 27, and 71 (*KRT25, KRT27*, and *KRT71*, respectively) that are expressed in the inner root sheath of hair follicles (33, 34) were highly expressed in SSL-RNAs (Fig. 4). These results suggest that SSL-RNAs provide significant information regarding the granular layer of the epidermis and the inner root sheath of hair follicles.

**Figure 4.**
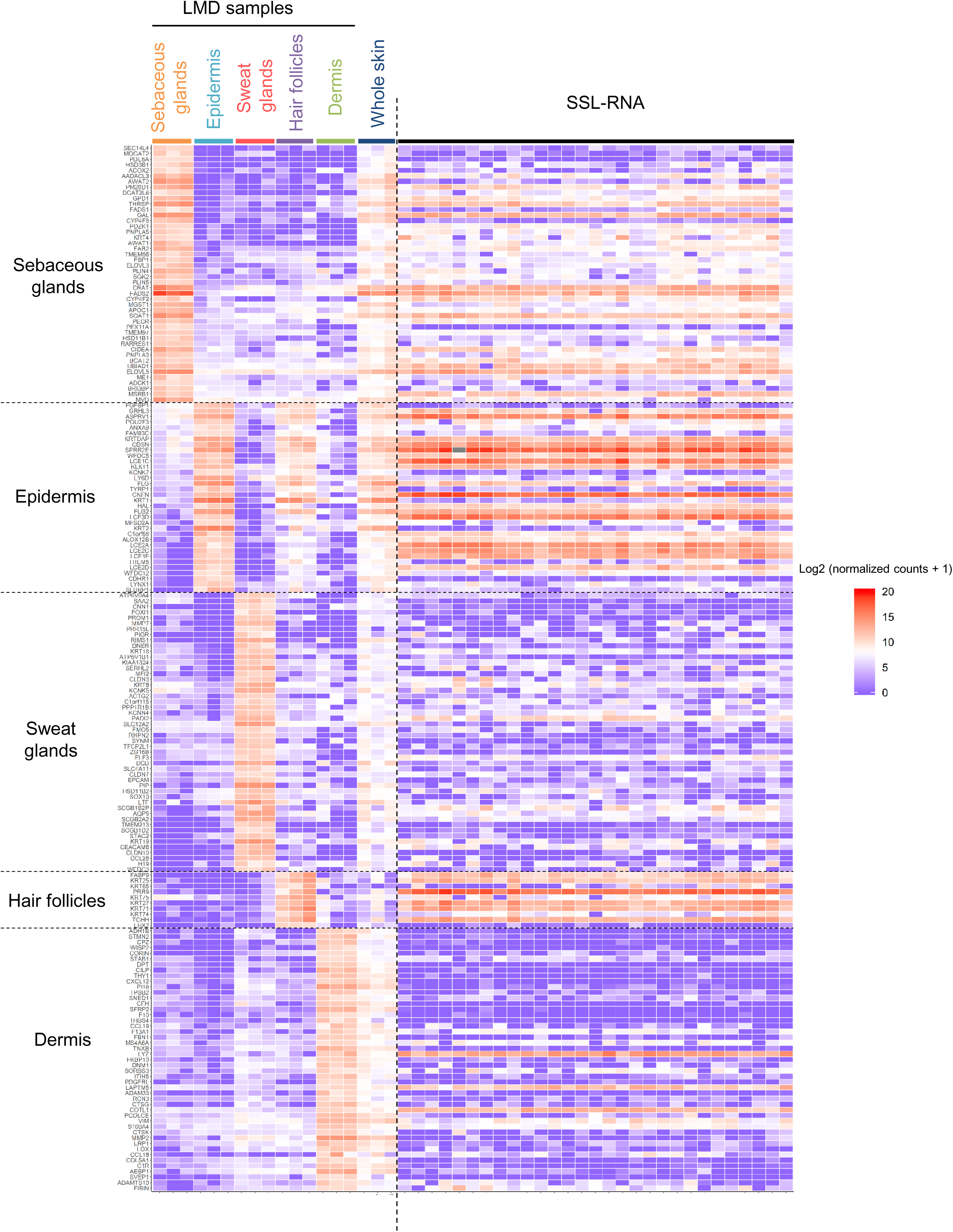
mRNA expression characteristic of different skin regions and its comparison with the expression profile of SSL-RNAs Heatmap showing the expression profiles for each region (sebaceous glands, epidermis, sweat glands, hair follicles, and dermis) isolated using LMD, whole skin from three healthy male subjects, and SSL- RNAs obtained from 29 healthy male subjects.

### Comparison of SSL-RNAs expression between healthy subjects and patients with AD

We analyzed SSL-RNAs from 29 healthy subjects and 30 patients with AD. The major output of the AmpliSeq data obtained with our modified method was as follows: i) the average number of reads was 11,456,318 in healthy subjects and 11,137,677 in patients with AD; ii) the average mapping ratio was 84.0 % in healthy subjects and 92.4 % in patients with AD; and iii) the average of the ratio of target detected genes was 44.8 % in healthy subjects and 50.2 % in patients with AD. First, we verified the expression of genes with significant differential expression in AD. Consistent with previous reports (39–42), our results in SSL-RNAs analysis showed that the expression of *CCL17,* interleukin 1β (*IL1B*), interleukin 13 (*IL13*), and S100 calcium binding protein A9 (*S100A9*) significantly increased in patients with AD, but the expression of *FLG* and involucrin (*IVL*) was significantly decreased in the patients with AD compared to healthy subjects (Fig. 5a). In a previous study reporting a global gene expression analysis in skin biopsy samples, genes related to the terminal differentiation of keratinocytes were significantly downregulated and those related to immune-mediated inflammation were upregulated in patients with AD compared with their expression levels in healthy subjects (39). Based on this report, we selected 12 genes related to terminal differentiation and 22 genes related to immune-mediated inflammation that were detected in SSL-RNAs and compared their expression patterns. Our results showed that the expression patterns of these genes in SSL- RNAs of patients with AD and healthy subjects were largely consistent with the previous report (Fig. 5b).

**Figure 5.**
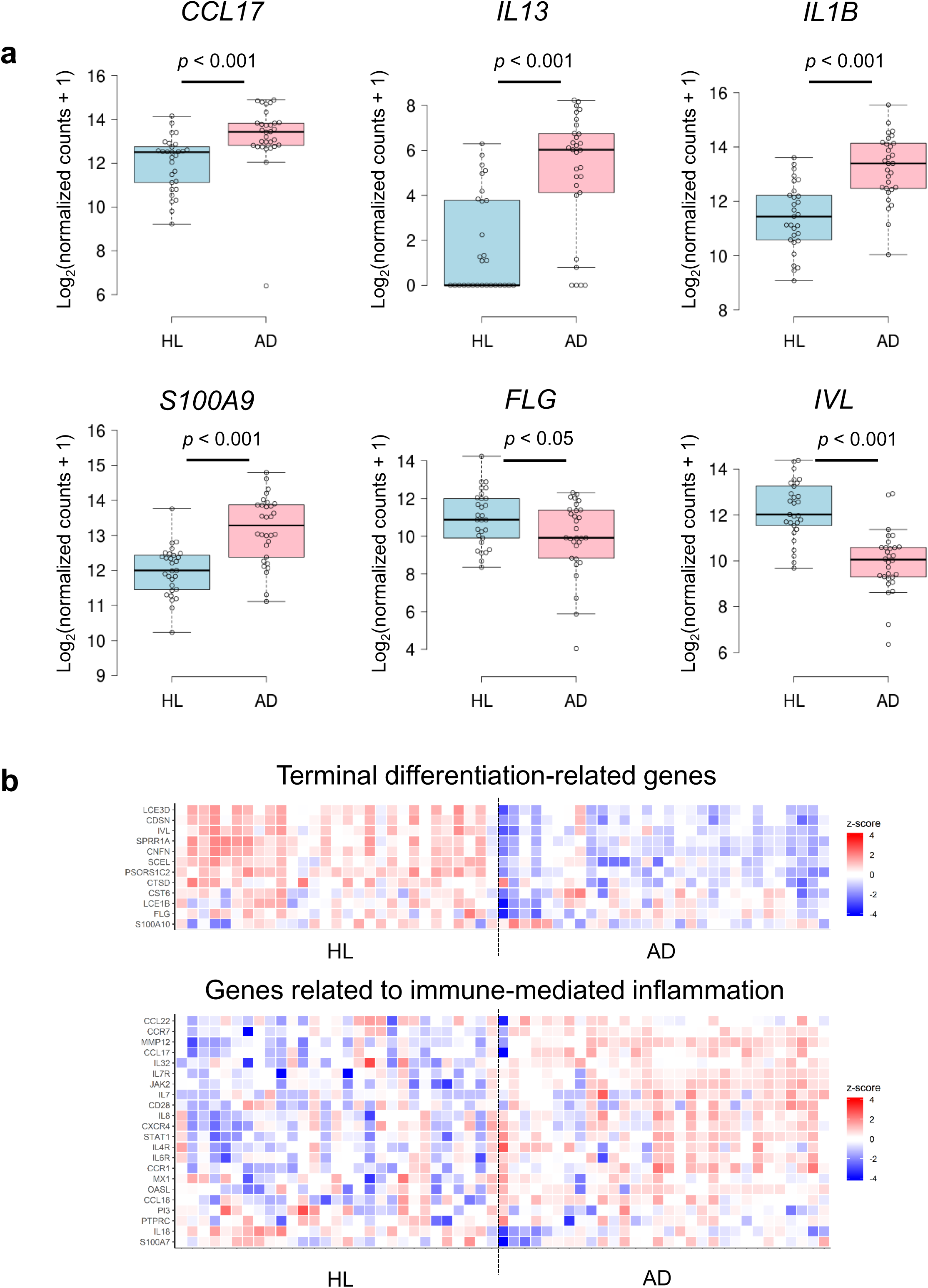
Comparison of AD marker genes in healthy subjects (HL) and patients with AD **(a)** The differential expression of *CCL17*, *IL13*, *IL1B, S100A9, FLG*, and *IVL* in HL and AD. Boxes represent mean ± interquartile range (IQR), and whiskers represent 1st and 3rd quartile 1.5 * IQR. Benjamini-Hochberg adjusted *p*-values are shown from the likelihood ratio test between HL and AD. HL (n = 29), AD (n = 30). **(b)** Heatmaps using z-transformed log2 (normalized counts + 1) in 12 terminal differentiation-related genes (upper) and 22 genes related to immune-mediated inflammation (lower).

Moreover, the analysis of the dimensionality reduction using t-distributed stochastic neighbor embedding (t-SNE) and variance stabilizing transformation (VST) values in all genes showed that the healthy subjects and patients with AD could be distinctly classified into two groups (Fig. 6a). To identify the differential biological functions between healthy subjects and patients with AD, we extracted 918 upregulated and 1,033 downregulated genes in patients with AD (Fig. 6b). In the 833 upregulated genes, GO terms of “mRNA splicing” and “stimulatory C-type lectin receptor signaling pathway” were significantly enriched, while GO term “detection of chemical stimulus involved in sensory perception of smell and keratinocyte differentiation” was enriched in the 951 downregulated genes (Fig. 6c).

**Figure 6.**
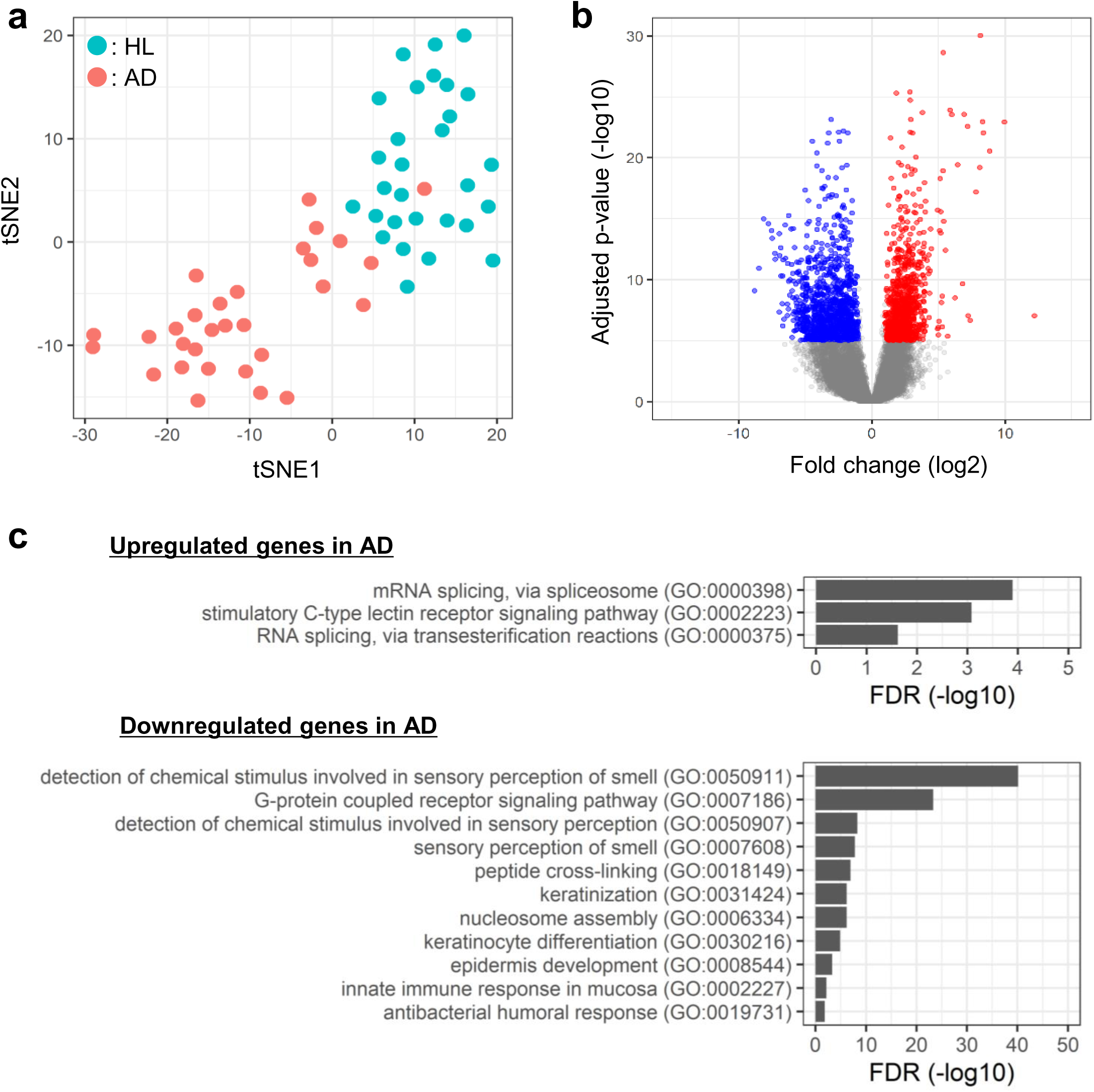
Characterization of SSL-RNAs profiles in healthy subjects (HL) and patients with AD **(a)** t-SNE analysis using variance stabilizing transformation (VST) values for all genes (green, HL; red, AD). **(b)** Volcano plot of differentially expressed genes (DEGs) (red, upregulated; blue, downregulated) in patients with AD compared to HL subjects (Benjamini-Hochberg adjusted *p*-value < 10^-5^ and fold change > 2.0). **(c)** Gene ontology analysis of DEGs. The upper panel shows significant biological process (BP) of upregulated DEGs and the lower panel shows BP of downregulated DEGs in patients with AD (FDR < 0.05).

The sebum secretion is reduced in patients with AD than in healthy individuals (43); however, the molecular mechanism underlying this reduction remains unknown. GO analysis was performed on 46 genes highly expressed in the sebaceous glands selected from the results of the LMD experiment (Fig. 4), which resulted in the enrichment of genes involved in lipid metabolism (Fig. 7a). The expression of 25 genes involved in lipid metabolism (GO:0006629) was downregulated in patients with AD compared to healthy individuals (Fig. 7b). Furthermore, genes encoding peroxisome proliferator-activated receptor alpha (*PPARA*), peroxisome proliferator-activated receptor gamma (*PPARG*), MYC proto-oncogene, bHLH transcription factor (*MYC*), transforming growth factor beta 1 (*TGFB1*), tumor protein p53 (*TP53*), and PR/SET domain 1 (*BLIMP1*) regulate sebocyte differentiation and sebum production *in vivo* and *ex vivo* (44–47). Among these gene, the expression of *TGFB1* was significantly upregulated in patients with AD than in healthy subjects (Fig. 7c).

**Figure 7.**
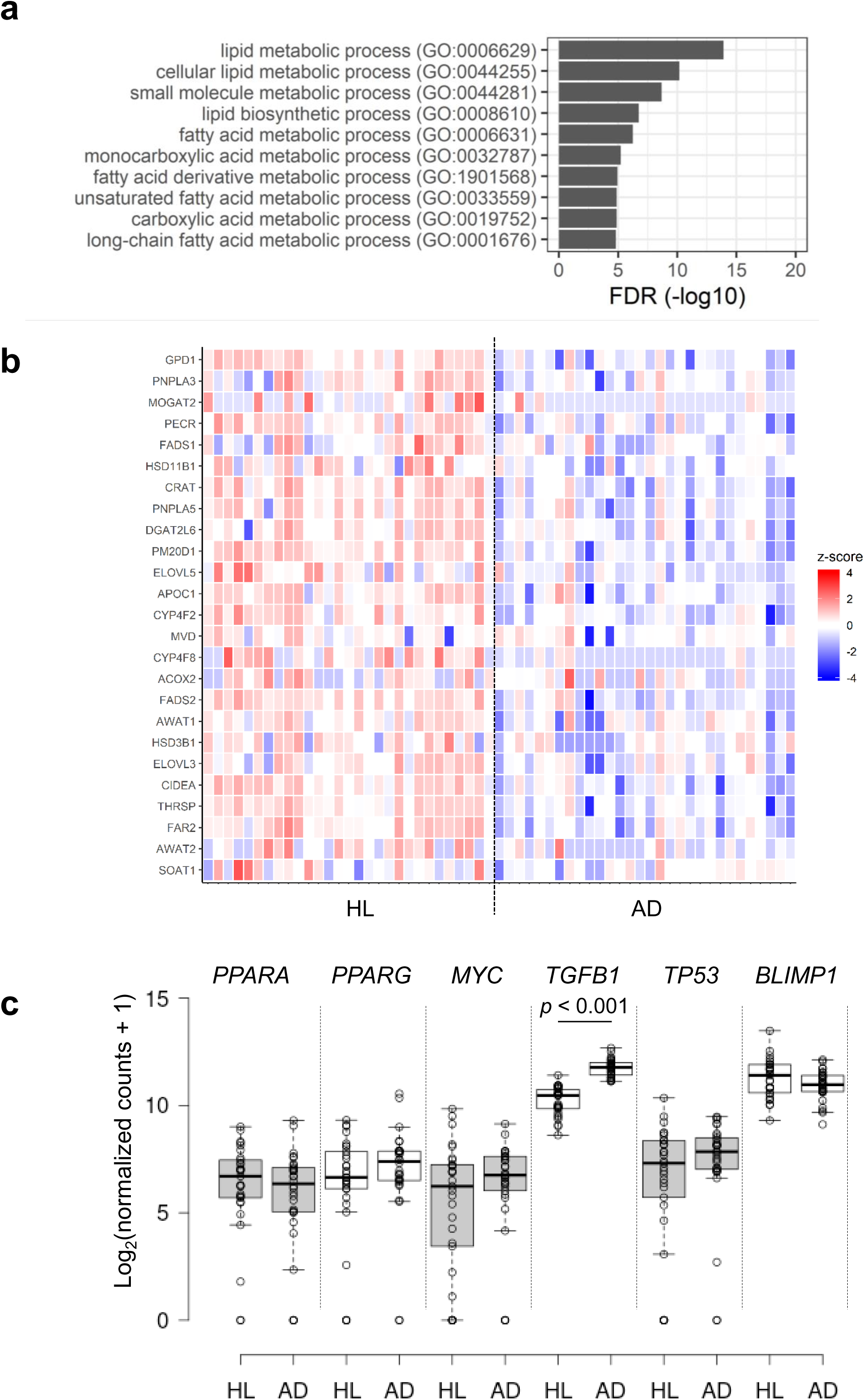
Comparison of SSL-RNAs profile representing highly expressed genes in sebaceous glands in healthy subjects (HL) and patients with AD **(a)** Gene ontology analysis of 25 genes highly expressed in sebaceous glands (selected in Fig. 4). **(b)** Heatmaps using z-transformed log2 (normalized counts + 1) of 25 genes highly expressed in sebaceous glands. **(c)** The differential expression of *PPARA, PPARG, MYC, TGFB1, TP53,* and *BLIMP1* in HL and patients with AD. Boxes represent mean ± interquartile range (IQR), and whiskers represent 1st and 3rd quartile 1.5 * IQR. Benjamini-Hochberg adjusted *p*-values are shown from the likelihood ratio test between HL and AD. HL (n = 29), patients with AD (n = 30).

## Discussion

In this study, we found that mRNAs of measurable quantity and quality were present in skin surface lipids. Further, for the first time, we established a non-invasive and comprehensive method to profile skin mRNAs using SSLs conveniently collected from the skin surface with an oil blotting film.

A significant quantity of RNases is present on the skin surface, and extreme caution should be taken when handling mRNAs. However, unexpectedly, we found that mRNAs in SSLs escape RNase degradation due to the lipid components of the sebum and can be analyzed by AmpliSeq transcriptome sequencing. We analyzed the lipid components responsible for inhibiting the RNase activity and our results suggested that FFAs, triacylglycerol, and squalene contributed significantly to the RNase inhibitory activity of sebum. Due to inhibitory effects of lipids on RNases, we speculate that mRNAs may be less susceptible to RNases-mediated degradation in the lipid-rich/low-water environment of SSLs. In addition, the optimal pH for RNase activity is 6.5-8.0 (48), whereas the skin surface is generally weakly acidic (pH 4.1 to 5.8) due to the presence of organic acids such as lactic acid (49); hence, these factors may collectively reduce the RNase activity in SSLs. It is known that sebum is secreted by sebaceous glands in the form of fine granules (4–5 nm) (50); however, there is no knowledge on the spatial arrangement of lipids, organic acids, and mRNAs in SSLs. Identifying the distribution of these components, as well as the molecular interactions between them, is necessary to understand the precise stabilization mechanism of mRNA on the skin.

We established the method for comprehensive analysis of SSL-RNAs. Recent advances in sequence technology have made it possible to analyze even degraded mRNA. To prepare RNA-seq libraries, a minimum DV200 value of 30 % is generally recommended. Using our modified method, the DV200 of human mRNAs in SSLs was approximately 56.5 %, making it suitable for transcriptome analysis. However, when the sequence libraries were prepared according to the standard protocol of Ion AmpliSeq Transcriptome Human Gene Expression Kit, the success rate was very low. Use of the optimized protocol (by standardizing conditions of reverse transcription and target amplification, and adding a purification step after target amplification) led to significant improvement in library production efficiency, and the transcriptome sequencing resulted in a success rate of 95 %. Further, the analytical error of this method was low and the results showed high correlation with qPCR results. Thus, our SSL-RNA analysis method using the improved AmpliSeq protocol enables profiling of the mRNA expression in SSLs in a reliable manner.

We investigated the origin of the SSL-RNAs by comparing their mRNA expression profile with those of different regions of the skin. Holocrine secretion of sebum made us speculate that the expression pattern of SSL-RNAs should be similar to that of the sebaceous glands. Indeed, analysis of different regions of the skin obtained by LMD showed that the mRNAs derived from the sebaceous glands were highly expressed in SSL-RNAs. However, SSL-RNAs were rich not only in mRNAs derived from sebaceous glands but also in those derived from the epidermis and hair follicles, the tissues in close contact with the sebum. In contrast, the mRNAs characteristic of sweat glands and dermis, which are not in close contact with sebum, were absent in SSLs. The mechanism underlying transfer of epidermal mRNAs into SSLs remain unclear, and there could be several possible mechanisms. The mRNAs characteristic of the granular layer of the epidermis (transcribed from *FLG*, *FLG2* and *ASPRV1*) were highly expressed in SSL-RNAs. In addition, stratum corneum is reported to contain detectable amounts of mRNA (13–15). We speculate that the epidermal mRNAs transferred to the surface of the stratum corneum due to keratinization are mixed with the sebum on the skin surface, leading to their presence in SSLs. The SSLs also contained hair follicle-derived RNAs, which may be related to the anatomical features of the hair follicle. SSL-RNAs were rich in mRNAs of *KRT25*, *KRT27*, and *KRT71*, the marker genes for the inner root sheath (33, 34). The inner root sheath detaches from the hair shaft and degrades during hair growth, and the process occurs at the orifice of the sebaceous duct (51). These observations indicate that the epithelial cells of the inner root sheath may get mixed with sebum at the orifice and skin surface, and as a result, information pertaining to the hair follicles may be reflected in the SSL-RNAs. In addition, since extracellular vesicles released from various cells contain several biomolecules including mRNAs (52, 53), it is possible that extracellular vesicles-derived mRNAs are also included in the SSL-RNAs. Collectively, our results indicate that SSL-RNAs predominantly contains mRNAs derived from the sebaceous glands, epidermis, and hair follicles, and are therefore, a useful resource for analyzing the biological information related to the relevant regions of the skin.

Finally, we verified the applicability of this method by performing a comparative analysis of the SSL-RNAs profiles of healthy subjects and patients with AD. We observed that the transcriptome profile was markedly different between healthy subjects and patients with AD, with differential expression of immune-mediated inflammation and terminal differentiation-related genes, as shown in a previous skin biopsy report (39). Moreover, the GO term “detection of chemical stimulus involved in sensory perception of smell” identified in our study was consistent with a previous report based on patients with AD (13) indicating that the analysis of SSL-RNAs successfully captured the characteristics of AD. Atrophy of the sebaceous glands and reduction in sebum secretion have been reported in patients with AD (54). However, little is known about the underlying mechanism, including the gene expression profile of sebaceous glands, in patients with AD due to difficulty in obtaining facial skin tissue samples containing sebaceous glands. Here, we showed that the expressions of 25 lipid metabolism-related genes highly expressed in sebaceous glands were lower in AD patients than in healthy subjects. Moreover, the expression of *TGFB1*, which suppresses sebocyte differentiation and lipid accumulation (47), was significantly increased in patients with AD, suggesting that the suppression of lipid synthesis via *TGFB1* may be one of the mechanisms responsible for dysregulated sebum synthesis in these patients. Thus, the transcriptome analysis of SSL-RNAs can evaluate the molecular profile of AD in a non-invasive manner, and is a promising method for comprehensive understanding of AD pathology.

In summary, we established a non-invasive method for SSL-RNA analysis that utilizes SSL samples collected by simply wiping the skin surface for less than a minute. This non-invasive method has potential application in unraveling the molecular profile of skin diseases, such as AD, which will be helpful for clinical management of these diseases in future. Understanding the status and course of AD at the molecular level is essential not only to assess the pathophysiology of AD but also to design its effective therapeutic treatments. The clinical phenotypes of AD are extremely complex, warranting the need for identifying biomarkers that can classify invisible endophenotypes (55). Further, since the skin is called “the disease-sensor organ” (56) and is believed to reflect the conditions inside the body, the SSL-RNA analysis may have wide applicability to understand various pathologies of human body.

### Data availability

The datasets generated and analyzed in the current study are available from the corresponding author on reasonable request.

## Author Contributions

T.I. conceived the study. T.I., A.H., Y.T., and T.M. planned the study. T.I., T.K., Y.U., M.Y., and N.Oy. performed the experiments and analyzed the data. T.M., Y.T., and N.Ot. supervised the research. T.I. and T.M. wrote the manuscript. All authors reviewed the manuscript.

