## Supplemental method 1 and figure 1 for "Non-invasive human skin transcriptome analysis using mRNA in skin surface lipids"

### **Supplementary Method 1:** Detailed protocol for the SSL-RNA analysis

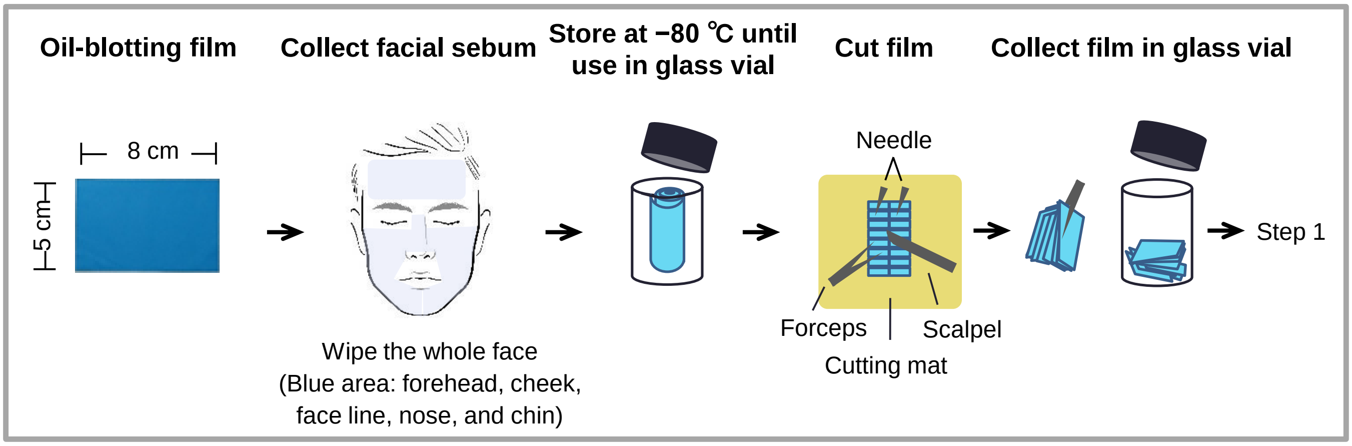


**Extraction of RNA from facial skin surface lipids (SSLs)**

1) Add 2.85 mL of QIAzol reagent to the glass vial, vortex, and transfer only the QIAzol solution to two fresh tubes.
2) Add 260 µL of chloroform to each tube, then vortex and centrifuge at 12,000 x *g* for 15 min at 4 ℃.

**Purification and concentration of SSL-RNA**

3) Transfer the upper layer (aqueous phase) to a fresh tube.
4) Purify total RNA using the RNeasy Mini kit (performing DNase treatment in the purification step) according to the manufacturer's instructions and elute from the resin twice with nuclease-free water (25 µL each time). Combine the eluates (100 µL each) into one tube (total 200 µL).
5) Concentrate total RNA by ethanol precipitation and finally dissolve in 10 µL of nuclease-free water.

**Library preparation with AmpliSeq**

6) Mix 1.75 µL of RNA solution with 0.5 µL of VILO reaction mix and 0.25 µL of SuperScript III Enzyme and perform reverse transcription under the following conditions: 25 ℃ for 10 min, 42 ℃ for 90 min, and then 85 ℃ for 5 min.
7) Mix 2.5 µL of cDNA solution, 1.5 µL of nuclease-free water, 2 µL of Ion AmpliSeq HiFi Mix, and 4.0 µL of Ion AmpliSeq Transcriptome Human Gene Expression Core Panel and amplify under the following conditions: 99 ℃ for 15 sec, 62 ℃ for 16 min, total 20 cycles (target amplification).
8) Purify the amplified DNA library with 10 µL of AMPure XP beads according to the manufacturer's instructions and then elute with 10 µL of nuclease-free water.
9) Check the quality of the DNA library with High Sensitivity D1000 ScreenTape and Agilent 4200 TapeStation. If the DNA library is amplified, a major band of about 170 bp will be observed.
10) Mix 3.5 µL of purified DNA library solution, 2 µL of Ion AmpliSeq HiFi Mix, 2 µL of Ion AmpliSeq Transcriptome Human Gene Expression Core Panel, 0.5 µL of VILO reaction mix, and 1 µL of FuPa. Incubate the tube under the following conditions: 50 ℃ for 10 min, 55 ℃ for 10 min, and 60 ℃ for 20 min to partially digest the primer sequences.
11) Add 2 µL of Switch solution, 1 µL of Ion Xpress Barcode Adapters, and 1 µL of DNA Ligase to 11 µL of reaction solution, followed by incubation at 22 ℃ for 60 min and 72 ℃ for 5 min to ligate the adaptor sequences.
12) Purify the DNA library with 18 µL of AMPure XP beads and then elute and dissolve using 50 µL of Library Amp Mix.
13) After adding 2 µL of Library Amp Primers, perform library amplification under the following conditions: 98 ℃ for 15 s, 64 ℃ for 1 min, total 5 cycles.
14) Purify the amplified DNA library with 25 µL of AMPure XP beads and then transfer the supernatant to a fresh PCR tube. Further, purify the library again with 60 µL of AMPure XP beads and then elute with 10 µL of TE.
15) Check the quality of the DNA library with High Sensitivity D1000 ScreenTape and Agilent 4200 TapeStation. If the DNA library is amplified, a major band of about 230 bp will be observed.

**RNA-seq**

16) Quantify the DNA library concentration using the Ion Library TaqMan Quantitation Kit.
17) After setting 50 pM DNA library in the Ion-chef system, perform emulsion PCR, template preparation, and chip loading.
18) Perform RNA-seq on the Ion-S5/X system.

**Supplementary Figure 1**. Similarity in mRNA expression profiles of SSL-RNA and each skin region.

(a) Tissue images of human sebaceous glands, epidermis, sweat glands, hair follicles, and dermis before and after LMD. Bar: 200 μm. (b) MDS analysis using the expression profile of each region and whole skin, n = 3. (c) mRNA expression profile of LMD samples and whole skin. The heatmap shows z-transformed expression values of marker genes in each LMD sample and whole skin, n = 3. SSL, skin surface lipids; LMD, laser microdissection

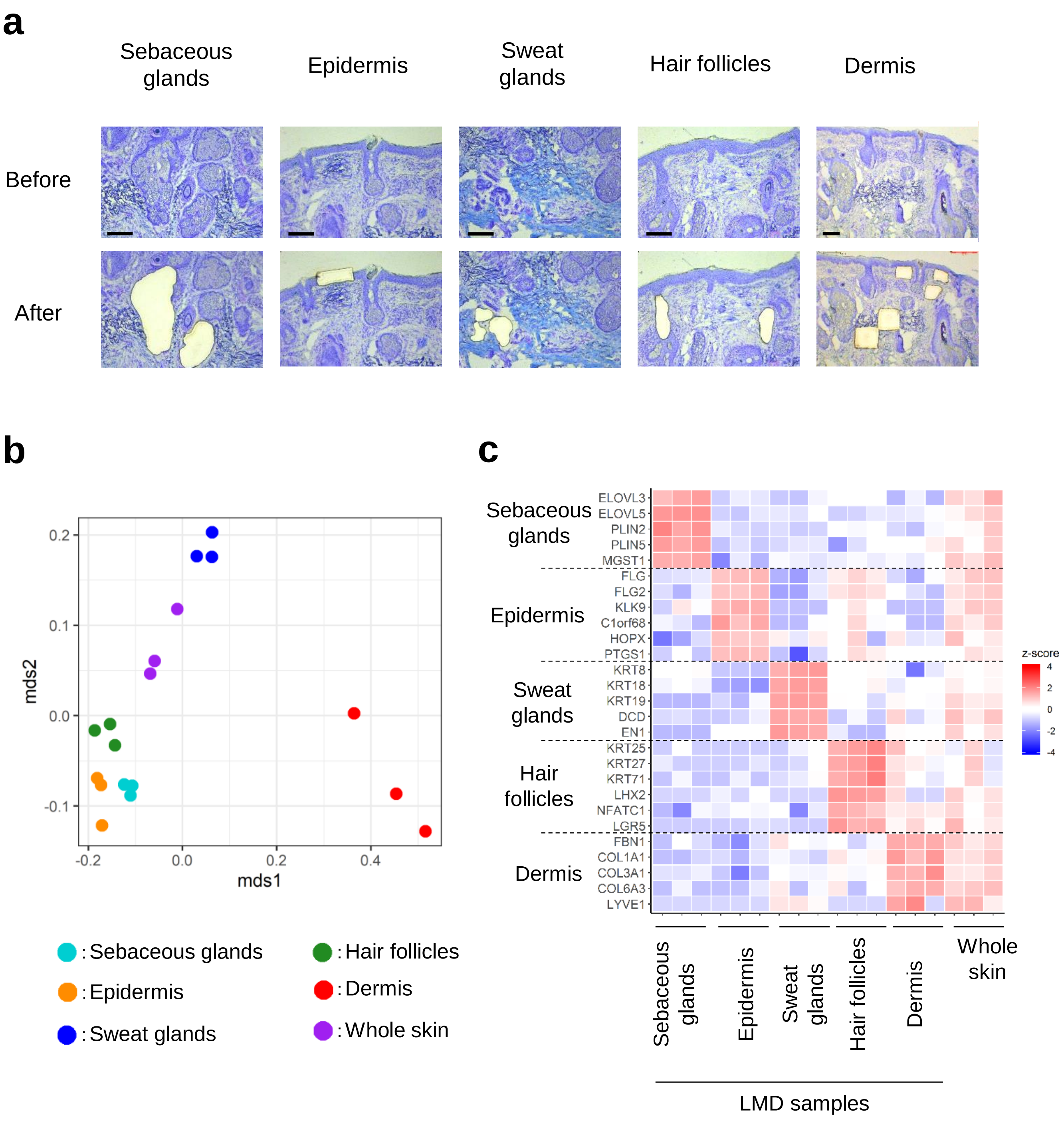
